# Widespread Corticothalamic Connectivity Identifies the Inferior Pulvinar as a Central Node in Visual Network Architecture

**DOI:** 10.64898/2026.03.02.709198

**Authors:** William C. Kwan, Angela Y. Fan, Andrea J Romanowski, Inaki-Carril Mundinano, Mitchell J. de Souza, James A. Bourne

## Abstract

The medial subdivision of the inferior pulvinar (PIm) has been implicated in motion processing, visuomotor integration, and residual visual function, yet a comprehensive account of its cortical inputs remains unresolved. Previous studies often relied on indirect cortical injections or tracer deposits spanning multiple pulvinar subdivisions, limiting anatomical specificity. Here, we used MRI-guided, cytoarchitectonically restricted retrograde tracer injections to selectively target PI in the common marmoset (*Callithrix jacchus*) and systematically map its cortical afferents.

Across four cases, retrogradely labeled neurons were widely distributed throughout occipital, temporal, parietal, and cingulate cortices, with a strong predominance in layer V, consistent with driver-like corticothalamic projections. Early and middle-tier visual areas (V1, V2, V3, V3A, V4, V6/DM) contributed substantial input, with labeling patterns corresponding to peripheral visual field representations. The middle temporal complex (MT, MTc, MST, FST) represented one of the densest sources of cortical projections. Prominent inputs also arose from posterior parietal regions, including LIP, MIP, VIP, AIP, and inferior parietal areas (e.g., PFG, OPt), linking PIm to visuospatial and action-related networks. Semi-quantitative analyses indicated that occipital cortex and the MT complex together accounted for approximately 60% of total cortical input, while parietal cortex contributed roughly 20%. Additional projections from retrosplenial and posterior cingulate cortices were observed.

These findings identify PIm as a central integrative node embedded within distributed visual and visuomotor networks. Rather than functioning as a restricted visual relay, PIm appears positioned to coordinate motion, spatial, and action-relevant signals within cortico-thalamocortical circuits supporting adaptive visually-guided behavior.

## 1 Introduction

Thalamocortical projections play a critical role in the transmission of sensory inputs from the periphery to the cortex. Within the visual sensory modality, the lateral geniculate nucleus (LGN) serves as the primary visual thalamic nucleus. Given the substantial connectivity between the retina and the primary visual cortex (V1), the body of literature that investigates thalamic control and modulation of vision is LGN-centric. Another visual thalamic nucleus of note is the pulvinar. Despite being the largest thalamic nuclei in anthropoids (Ohye, 2002), the pulvinar has attracted significantly less investigation, which could be owed to the fact that retinal afferents within the pulvinar are less prolific when compared to the LGN and even the superior colliculus within the midbrain (Kwan et al., 2019). Whilst the pulvinar is somewhat enigmatic, the immense body of work which profiled the cytoarchitecture of the pulvinar (see Baldwin and Bourne, 2020 for a comprehensive review) and mapped pulvino-cortical circuits through neural tracers (Kaas and Lyon, 2007) has provided invaluable insights into its functional role in vision.

The pulvinar is divided into three major subdivisions: the medial (PM), lateral (PL), and inferior (PI) pulvinar. PM, the largest division, is interconnected with cortical areas spanning multiple sensory modalities and with frontal regions involved in cognitive and executive functions (Homman-Ludiye et al., 2019; Córdoba-Claros et al., 2025). In contrast, PL and PI project predominantly to visual cortical areas (Kaas and Lyon, 2007). PI can be further subdivided into four subnuclei: posterior (PIp), medial (PIm), central medial (PIcm), and central lateral (PIcl) **(see Fig. 1C)**.

**Figure 1:**
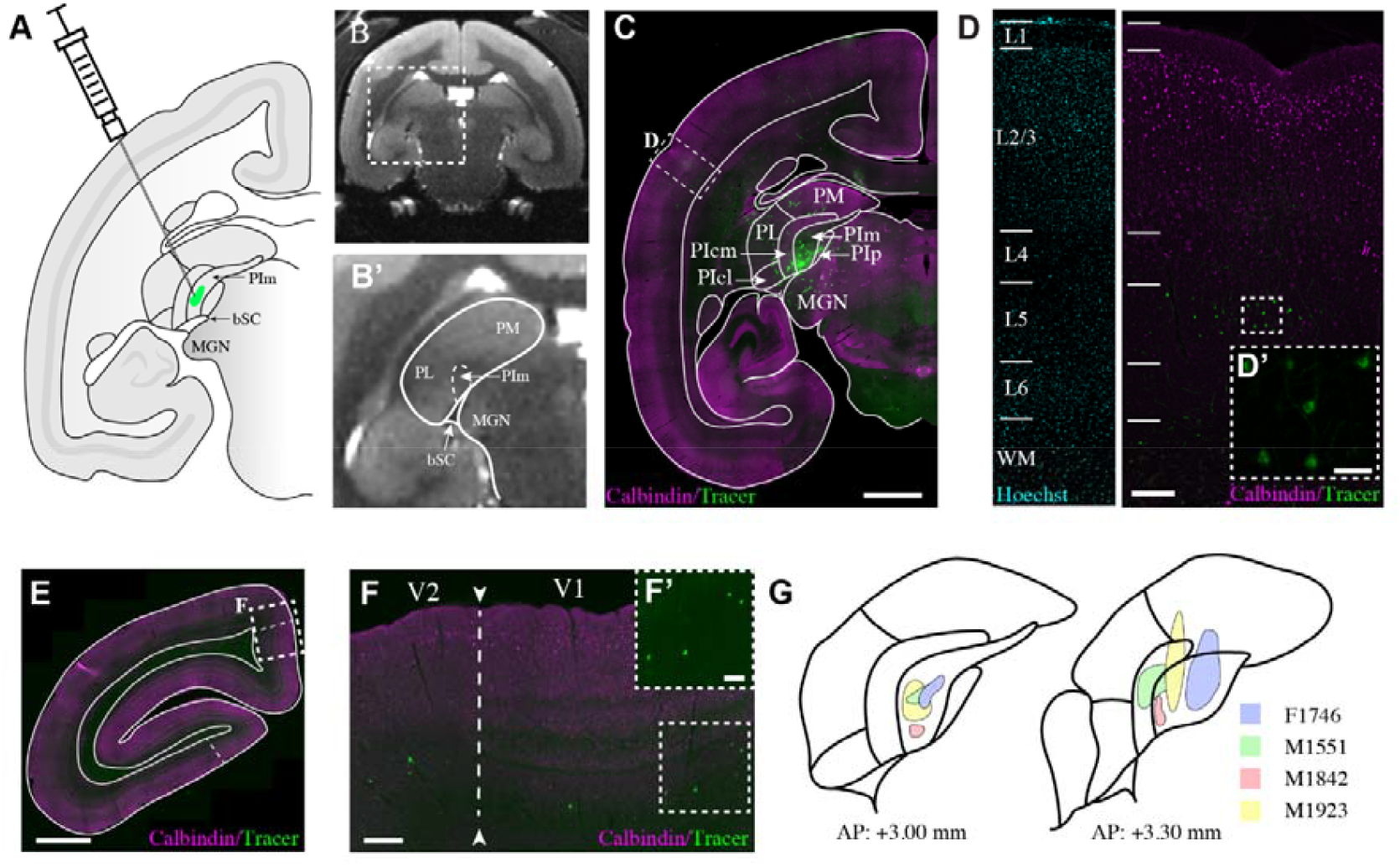
Mapping cortical inputs to the medial subdivision of the inferior pulvinar (PIm) with retrograde neural tracer injections. **A**. Schematic of the marmoset brain, the pulvinar nucleus, and PIm in coronal view. **B**. T2 MRI of the marmoset brain. **B’**. Zoomed-in image of pulvinar from **B. C**. Low-powered photomicrograph of case M1842 following tracer deposition in the temporal cortex at coordinate AP: +3.00mm. scale bar = 2mm **D**. Retrograde label is largely restricted to layer 5, as evident in **D’**. The scale bars in the low- and high-powered photomicrographs are 500 µm and 50 µm, respectively. **E**. Low-powered photomicrograph of case M1842 following tracer deposition in the occipital cortex at coordinate AP: -5.00 mm. Scale bar = 2 mm. **F**. Higher powered photomicrograph of the visual cortex as outlined in E. As evident in **F’**, V1 and V2 projections to PIm are localised in infragranular layers. Scale bar = 500 µm and 100 µm in F and F’, respectively. **G**. Summary of the location of the tracer bolus in each subject.

A defining feature of PI is its receipt of direct retinal input, with PIm serving as the principal locus of retinal terminations (O’Brien et al., 2001; Warner et al., 2010). PIm is distinguished by the absence of calbindin immunoreactivity and is bordered by calbindin-rich PIcm and PIp (Cusick et al., 1993; Stepniewska and Kaas, 1997). Notably, PIm contains a complete retinotopic representation of the visual field (Mundinano et al., 2019; Arafune-Mishima et al., 2020). Taken together, this evidence supports the proposal that PIm may function as a first-order visual thalamic relay analogous to the LGN (Sherman and Guillery, 1998).

PIm projects to multiple cortical areas (Kaas and Lyon, 2007), with its projection to the middle temporal area (MT) being particularly well characterized (Warner et al., 2010). MT develops in parallel with primary sensory cortices (Bourne and Rosa, 2005) and plays a central role in motion processing (Born and Bradley, 2005). The PIm–MT pathway has attracted considerable interest as a potential substrate for residual visual function following V1 lesions, commonly referred to as blindsight (Warner et al., 2010; Mundinano et al., 2018; Kinoshita et al., 2019; Fox et al., 2020; Takakuwa et al., 2021).

Beyond its proposed role in blindsight, the broader contribution of PIm to visual processing remains unclear. PIm receives input predominantly from wide-field retinal ganglion cells, with minimal contribution from midget and parasol cells that dominate LGN projections (Grünert et al., 2021). Subcortically, PIm connectivity is relatively restricted, forming connections with other retinorecipient nuclei including the parabigeminal nucleus (Diamond et al., 1992), pregeniculate, and subgeniculate nuclei (Kwan et al., 2019). In contrast, projections from the superior colliculus and LGN are sparse in PIm but dense in adjacent PIcm and PIp (Kwan et al., 2019).

Electrophysiological recordings demonstrate that PIm neurons respond to moving stimuli, with a subset receiving descending input from MT exhibiting direction selectivity (Berman and Wurtz, 2011). Functional imaging further indicates responsiveness to looming stimuli (Cléry et al., 2020). Lesion studies suggest that disruption of PIm impairs visuomotor development (Mundinano et al., 2018) and visual attention to target objects (Snow et al., 2009; Wilke et al., 2010).

Because the literature concerning the functional organization of the PIm remains comparatively fragmented, we were motivated to generate a cortical connectivity map centered specifically on this subdivision of the inferior pulvinar. Establishing a detailed and anatomically precise framework of PIm afferent connectivity is a necessary first step toward formulating targeted hypotheses about its circuit-level contributions to visual and visuomotor processing. Although neuroanatomical tracing studies examining pulvinar connectivity are not unprecedented, most prior investigations have inferred PIm connectivity indirectly by placing retrograde tracers in cortical target regions and subsequently identifying labeled neurons within PIm (Kaas and Lyon, 2007; Gattass et al., 2018; Huo et al., 2019). While highly informative, this strategy does not systematically resolve the full complement of cortical inputs to PIm, nor does it permit a comprehensive evaluation of the relative strength or topographic organization of these projections. Moreover, earlier surveys that directly injected tracers into the inferior pulvinar frequently relied on relatively large tracer deposits that extended across multiple pulvinar subdivisions, thereby limiting the anatomical specificity of the resulting connectivity maps (Benevento and Rezak, 1976; Ogren and Hendrickson, 1977; Dick et al., 1991; Marion et al., 2013). Given the cytoarchitectonic and connectional heterogeneity of pulvinar subdivisions, such encroachment likely obscured subdivision-specific connectivity patterns and contributed to inconsistencies across studies.

In the present study, we therefore confined neural tracer injections predominantly to cytoarchitectonically defined PIm in the marmoset monkey (*Callithrix jacchus*), enabling a more selective and subdivision-resolved assessment of its cortical afferents. Through a systematic and extensive survey of labeled cortical neurons, we confirm several previously described PIm– cortical pathways and identify additional cortical regions that, to our knowledge, have not been reported to project to PIm. These findings refine and expand the existing model of PIm connectivity, suggesting that PIm is embedded within a broader and more integrative cortical network than previously appreciated. By establishing a reliable and reproducible methodological framework for selectively targeting PIm circuits and obtaining functional readouts, this revised network map offers a foundational resource for future studies aimed at elucidating the circuit mechanisms and behavioral significance of PIm within primate visual systems.

## 2 Materials and Methods

### Animals

Four adult New World marmoset monkeys (*Callithrix jacchus*) (1.7 – 4.1 years of age) sourced from the National Nonhuman Primate Breeding and Research Facility (Australia) were used in this study. Animals received administration of neural tracers within PIm. Following surgical procedures, the animals underwent 7 days of recovery before euthanasia. All experiments were conducted in accordance with the Australian Code of Practice for the Care and Use of Animals for Scientific Purposes, and were approved by the Monash University Animal Ethics Committee, which also monitored the health and wellbeing of the animals throughout the experiments.

### MRI Guided Microinjections

All animals in this study underwent delivery of a neural tracers under MRI guidance. Full details of preparation of animals and visualisation of target areas have been described previously (Mundinano et al., 2016). In brief, animals are anaesthetized with an intramuscular injection of alfaxalone (8mg • kg-1) combined with diazepam (3mg • kg-1) with maintenance of anaesthesia achieved with inspired isoflurane (0.5% - 2.0% in medical oxygen at 500mL • min-1). Animals were additionally medicated with atropine sulfate (50 µg • kg-1) to increase cardiac muscle contraction and reduce mucosal secretions, and with dexamethasone (40 µg • kg-1) to minimise cerebral oedema. Once an appropriate plane of anaesthesia was achieved, animals were placed in a custom-built non-ferromagnetic stereotaxic frame to facilitate T2-weighted MR Imaging. Scans were exported as Digital Imaging and Communications in Medicine (DICOM) files, and the brain images were visualized using Horos version 3.3.5 software (https://horosproject.org) to delineate the boundaries of PIm or MT. Our target area PIm received 100 nl of neural tracer at a rate of 300 nl • min^-1^ through a Hamilton Neuros Syringe (Hamilton; Cat# 65460-03) fitted with customized 33G 15º bevelled needles (Hamilton; Cat# 65461-02). At the completion of the microinjection procedure, the cranium was reconstructed and repaired with bone wax and veterinary tissue adhesive (Vetbond) as required and the scalp was sutured.

### Neural Tracers

To label descending cortical inputs to PIm, we administered a neural tracers that consisted of a [1:1] cocktail of the retrograde tracer cholera toxin subunit B (CTB) (recombinant) with Alexa Fluor 488 [10mg • ml^-1^] (Thermo Fisher Scientific, Waltham, MA; Cat# C-22841), combined with the anterograde tracer dextran amine, Alexa Fluor 488; 10,000 MW [10% w/v] (Thermo Fisher Scientific; Cat# D22910).

### Histology and immunohistochemical tissue processing

When animals reached their endpoint (seven days post tracer delivery or at the conclusion of behavioural experimentation), they were humanely euthanized with an overdose of sodium pentobarbitone (>100mg/kg). When animals were unconscious as denoted by the absence of withdrawal and corneal reflex, animals were transcardially perfused with 10mM PBS that has been supplemented with heparin (50 IU/ml of PBS), followed by 4% paraformaldehyde in 10mM PBS. Brains were post-fixed in 4% paraformaldehyde then dehydrated in serial solutions of PBS-sucrose before being snap frozen in 2-methylbutane that has been chilled to -50ºC and stored in a -80ºC freezer until cryosectioning.

Brain samples were cryosectioned in the coronal plane at 50µm and stored free-floating in a cryoprotective solution (as outlined in previous work (Bourne and Rosa, 2005). Free-floating sections were labelled with the calcium-binding protein, calbindin (CB; Dilution: 1:3000; Swant cat# CB38; RRID: AB_10000340) and visualized with secondary antibody conjugated to Alexa Fluor 647 (Dilution: 1:1000; Thermo Fisher, Cat # A32795, RRID: AB_2762835) to delineate cortical and pulvinar boundaries. Sections were counterstained with Hoechst 33258 (Dilution 1:1000; Thermo Fisher, Cat # H3569; RRID: AB_2651133) to label cell nuclei and delineate cortical layers.

### Microscopy and digital image processing

Brain sections were imaged on a wide-field fluorescence microscope (Leica Thunder) using a 10x objective. Images were obtained using Leica LAS X software (Leica) at a resolution of 1.6 pixels/µm and saved in LIF format. Individual colour channels were split using FIJI (Schindelin et al., 2012) and exported as PNG. Image dimensions were scaled down to 30% of their original size with resampling to improve handling of image datasets within Adobe Illustrator. No digital adjustments were made to brightness and contrast for analysis and quantification of tracer label.

### Delineation of area boundaries and representation of tracer label

Brain section contours, boundaries, line art and digital representation of retrograde labelled cells were performed manually and drawn using Adobe Illustrator following import of fluorescently labelled microscopy images (1.6 pixels / µm). Delineation of area boundaries was undertaken by expert marmoset neuroanatomists (W.C.K, A.Y.F, I-C.M & J.A.B). We examined the laminar cytoarchitecture of Hoechst nuclear stain as well as calbindin immunolabel in the cortex for each animal and cross referenced this laminar profile with distinct anatomical landmarks such as lateral ventricle size/shape, relative position of large subcortical structures such as the SC or LGN, white matter tract size/shape.

We further cross referenced our calbindin and Hoechst label with the cortical area boundaries and nomenclature as defined in a marmoset brain stereotaxic atlas (Paxinos et al., 2012). The convergence of the labelling we observed from the calbindin and Hoechst labelling in our subjects and how areal boundaries are defined in the marmoset atlas permitted us to reach a consensus on the area boundaries within each brain slice in each of our subjects.

Following boundary delineation, cells were manually counted using the Cell Counter plugin in FIJI. For each animal, the number of PIm-projecting cells within each area was normalized to the total number of retrogradely labelled cells, yielding a semi-quantitative measure of the strength of connectivity between each cortical area and PIm.

## 3 Results

### 3.1 A primate model for characterizing PIm connectivity

To elucidate the connectivity of the medial inferior pulvinar (PIm), we mapped its projections in the common marmoset (*Callithrix jacchus*), a New World primate whose pulvinar organization closely parallels that of other primates, including humans (Münkle et al., 2000; Baldwin and Bourne, 2020), and in which the first emergence of PIm in primates. The marmoset’s small body size offers a distinct experimental advantage, enabling precise, MRI-guided subcortical microinjections (Mundinano et al., 2016) **(Fig. 1A)**. We targeted PIm in four adult animals and delivered neural tracers via pressure injection to define its connectional architecture. Across cases, retrogradely labelled neurons were predominantly confined to layer V **(Fig. 3.1C, D)**, revealing a striking laminar specificity. Injection sites were localized to the rostral-ventral extent of PIm **(Fig. 3.1G)**, corresponding to peripheral visual field representations consistent with known retinotopic specializations (Mundinano et al., 2019). In subject M1842 we were able to completely restrict the tracer deposit to PIm. In subject M1551 the tracer bolus encroached on the adjacent PIcm but was still within PI. In two cases (M1923 & F1746), limited tracer backflow extended into adjacent PI subdivisions and PM. Nevertheless, all four cases were analyzed collectively and the convergent patterns observed across our dataset was cross-referenced to case M1842 to derive conclusions regarding PIm connectivity.

### 3.2 Early as well as higher-order visual cortex projects to PIm

The flow of information within the visual cortex is typically hierarchical, reflecting the progressive transformation of retinal signals as they pass through successive synaptic relays. The primary visual cortex (V1) receives the majority of retinal afferents via the thalamus and serves as the base tier in the hierarchy before signals flow onward to higher cortical areas (Felleman and Van Essen, 1991). Although strict boundaries between hierarchical “tiers” remain debated, functional and anatomical frameworks generally distinguish early visual areas (including V1 and V2), intermediate extrastriate regions within occipital cortex (e.g., V3, V3A, V4, MT-complex subdivisions), and higher-order association areas in temporal and parietal cortex that support integrative visual processing (Felleman and Van Essen, 1991; Ungerleider and Haxby, 1994; Kosslyn and Thompson, 2003).

**Table 1:** Abbreviations of cortical areas in the marmoset monkey. Nomenclature used follows the marmoset brain stereotaxic atlas (Paxinos et al., 2012)

| Abbreviation | Full Name |
| --- | --- |
| A19DI | Area 19 of cortex, dorsointermediate part |
| A19M | Area 19 of cortex, medial part |
| A23a-b | Area 23a-b of cortex |
| A23V | Area 23 of cortex, ventral part |
| A29a-d | Area 29a-d of cortex |
| A30 | Area 30 of cortex |
| A31 | Area 31 of cortex |
| A35 | Area 35 of cortex |
| A36 | Area 36 of cortex |
| AuA1 | Auditory cortex, primary area |
| AuCL | Auditory cortex, caudolateral area |
| AuCM | Auditory cortex, caudomedial area |
| AuCPB | Auditory cortex, caudal parabelt area |
| AuML | Auditory cortex, middle lateral area |
| CA1 | Hippocampus field CA1 |
| FST | Fundus of superior temporal sulcus area of cortex |
| LIP | Lateral intraparietal area of the cortex |
| MIP | Medial intraparietal area of the cortex |
| MST | Middle superior temporal area of cortex |
| MT (V5) | Middle temporal area/visual area 5 |
| MTc | Middle Temporal Crescent |
| OPt | Occipito-parietal transitional area of the cortex |
| PE | Parietal area PE |
| PEC | Parietal area PE, caudal part |
| PF | Parietal area PF |
| PFG | Parietal area PFG |
| PG | Parietal PG, medial |
| FSTv (PGa-Ipa) | Fundus of superior temporal ventral area (Parietal Areas PGa and IPa) |
| PGM | Parietal area PG, medial part |
| ProSt | Prostriate area |
| TE2 | Temporal area TE2 (inferior temporal cortex) |
| TE3 | Temporal area TE3 (inferior temporal cortex) |
| TEO | Temporal area TE, occipital part |
| TF | Temporal area TF |
| TFO | Temporal area TF, occipital part |
| TH | Temporal area TH |
| TL | Temporal area TL |
| TLO | Temporal area TL, occipital part |
| TPt | Temporoparietal transitional area |
| V1 | Primary visual cortex |
| V2 | Visual area 2 |
| V3/VLP | Visual area 3/ventrolateral posterior area |
| V3A/DA | Visual area 3A/dorsoanterior area |
| V4/VLA | Visual area 4/ventrolateral anterior area |
| V6/DM | Visual area 6/dorsomedial area |
| V6A/PPM | Visual area 6A/posterior parietal medial area) |
| VIP | Ventral intraparietal area of cortex |

For the purposes of this study, we defined early-tier visual cortex as V1 and V2. Middle-tier areas included remaining occipital extrastriate regions (e.g., V3 and related subdivisions), while higher-order visual areas were defined as those within temporal and posterior parietal domains implicated in advanced perceptual and visuospatial computations.

Retrograde tracing revealed that PIm receives substantial input from both early and middle-tier visual areas. Labeled neurons were concentrated along the lateral and dorsal banks of occipital cortex, with a strong predominance of layer 5 projections arising from V1, V2, V3, V3A, A19DI, V4, and V6/DM (Fig. 2–5A–D). This laminar pattern is consistent with established corticothalamic projection motifs, in which layer 5 neurons provide driving input to higher-order thalamic nuclei (Sherman and Guillery, 2002), suggesting that PIm is embedded within a distributed occipital network spanning multiple hierarchical stages of visual processing.

**Figure 2:**
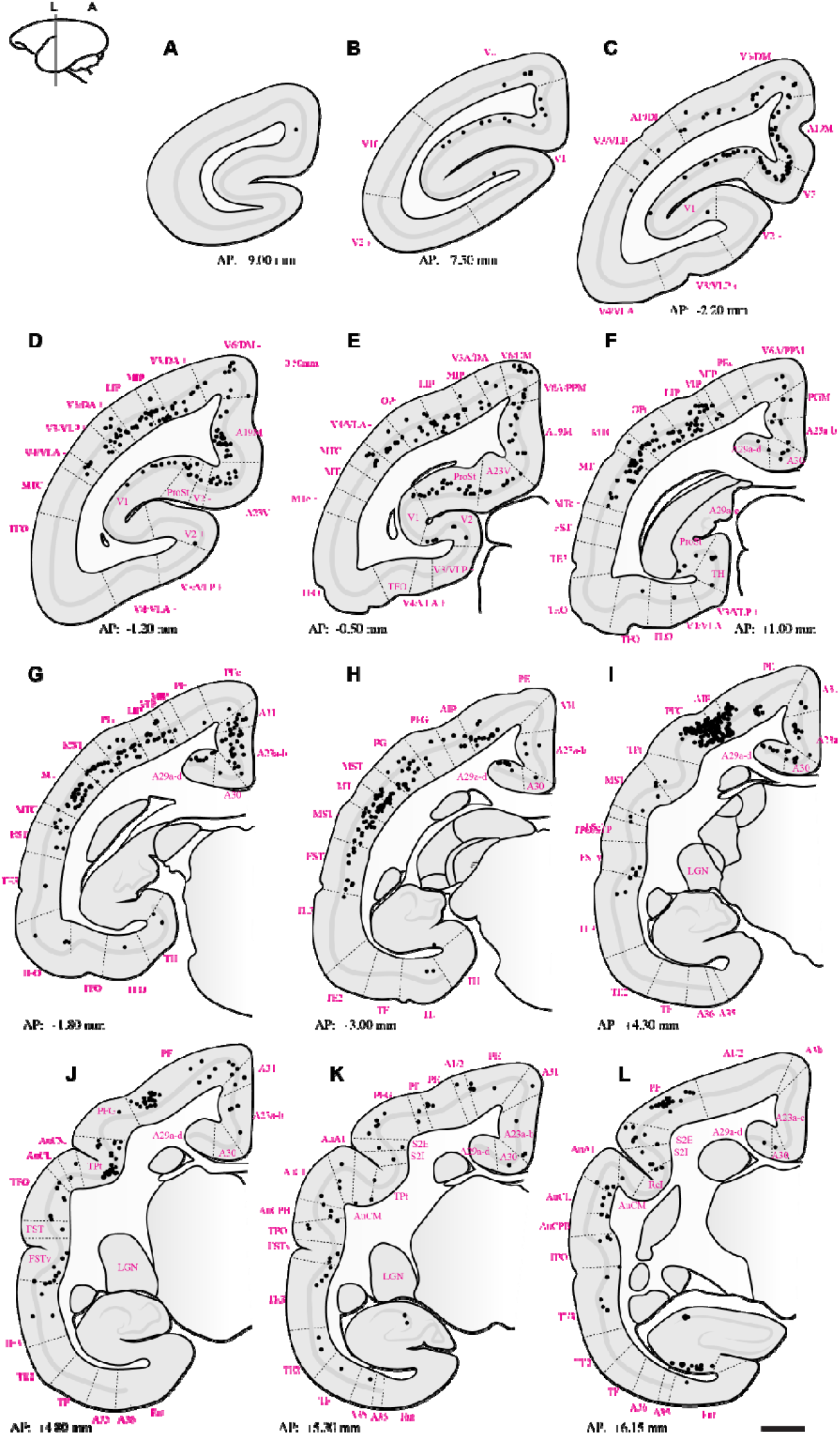
Summary of retrograde label throughout case **F1746**. Sections starting at **A**. are the most caudal, moving to the most rostral. Black dots indicate retrograde labelled cells following tracer deposition in PIm (refer to Fig 1G for tracer bolus placement). Dotted lines indicate boundaries of cortical areas. Abbreviations of cortical areas summarised in Table 1. Scale bar = 2mm.

Previous investigations of pulvinar afferent connectivity in marmoset and macaque revealed the existence of projections from early visual areas within PI (Ogren and Hendrickson, 1977; Kennedy and Bullier, 1985; Dick et al., 1991; Fitzgibbon et al., 2007; Baldwin et al., 2012) (Ogren and Hendrickson, 1976; Kennedy and Bullier, 1985; Dick et al., 1991; FitzGibbon et al., 2007; Baldwin et al., 2012). These early expeditions, which mapped visual cortical efferents in the pulvinar, only described which major subdivision received the cortical signals (e.g. the inferior pulvinar as a whole). Later studies, which examined PI connectivity, utilized calbindin immunohistochemistry to delineate PI into the four canonically described subdivisions and showed projections from early striate areas to PI primarily terminated in the subdivisions that flank PIm, i.e. PIcm, PIcl, and PL (Cusick et al., 1993; Kaas and Lyon, 2007; Gattass et al., 2018).

Whilst our results of V1/V2 projections to PIm have not been widely reported, they agree with a previous study which observed V1 neurons from peripheral visuotopic representations projecting to PIm (Gutierrez and Cusick, 1997). The foveal visual field in PIm is predominantly represented by its ‘dorsomedial tail’, whilst the ventral ‘base’ of PIm, which is where we deposited tracer in all four cases, is represented by the periphery (Mundinano et al., 2019). The V1/V2 retrograde label we observed in the calcarine sulcus in more rostral portions of V1 **(Fig. 2-5: A-C)** corresponds to the peripheral visual field (Fritsches and Rosa, 1996) and aligns with our placement of neural tracer. We were unable to deposit tracer into the foveal representation of PIm in this study. As such, we couldn’t definitively exclude the existence of foveal connectivity between PIm and the early visual cortex. However, prior work indicates that pulvinar connectivity associated with foveal representations is minimal or absent (Kaas and Lyon, 2007). On this basis, we infer that the visual input PIm receives from the early visual cortex is preferentially weighted toward peripheral visual field representations.

The pattern of labeling observed in higher-order visual cortical areas, including V3, V3A, A19DI, and V6/DM **(Fig. 2–5: C–D)**, is consistent with previously reported corticocortical connectivity patterns described in both New World and Old World primates (Asanuma et al., 1985; Dick et al., 1991; Yeterian and Pandya, 1997; Kaas and Lyon, 2007). These earlier tract-tracing studies demonstrated robust projections between extrastriate visual areas and posterior parietal regions, supporting the interpretation that the present labeling reflects homologous higher-order visual pathways.

To our knowledge, direct projections from V4/VLA to PIm have not previously been reported in the marmoset. In contrast, anatomical connectivity between V4 and subdivisions of the inferior pulvinar, including PIm, is well established in the macaque based on retrograde and anterograde tracer studies (Gattass et al., 2014). The present findings therefore extend existing data by identifying a comparable projection in the marmoset, suggesting that this thalamocortical network may be more evolutionarily conserved across primate lineages than previously recognized.

From regions occupying the medial surface of the extrastriate cortex and the calcarine sulcus, we observed retrogradely labelled neurons in areas A19M, A23V, and prostriata (ProSt) following retrograde tracer injections in PIm **(Fig. 2–5: D, E)**. Although pulvinar connectivity with these areas has not previously been reported in the marmoset, a study in the macaque demonstrated axonal terminal labelling in PIm following broad anterograde tracer injections in medial extrastriate cortex (Yeterian and Pandya, 1997). Functional investigations have revealed that these areas are preferentially involved in processing visual motion perceived from the far peripheral visual field (Palmer and Rosa, 2006; Yu et al., 2012; Mikellidou et al., 2017). Consistent with this functional specialization, retrograde labelling was observed in A19M and ProSt in all four cases, and in A23V in three cases. This pattern may reflect tracer uptake within the peripheral representation in ventral PIm.

### 3.3 Projections from the temporal and parietal cortices

As visual processing evolved in nonhuman primates and humans, the computations underlying visual perception and visually guided behaviour became increasingly complex, accompanied by an expansion of extrastriate cortex into temporal and parietal domains. Subcortical circuits play a critical role in mediating information transfer between major cortical lobes, with the pulvinar serving as a key hub for regulating communication across distributed visual cortical networks (Saalmann et al., 2012). Awake functional MRI studies in nonhuman primates demonstrate that visual stimulation engages widespread pulvinocortical networks, with activation patterns varying systematically according to the nature of the visual stimulus presented (Cléry et al., 2020; Guedj and Vuilleumier, 2023). Given the proposed integrative role of the pulvinar in large-scale cortical communication, we sought to determine whether tracer labelling in our cases similarly revealed extensive connectivity with parietal and temporal cortical regions.

Extensive retrograde labelling was observed in the temporal cortex **(Fig. 2–5: E–I)**, with area MT exhibiting the densest cortical projections to PIm. Substantial projections to PIm were also identified from MT satellite regions, including MTc, MST, and FST. The distribution of retrograde labelling within these motion-related temporal areas is consistent with previous tract-tracing studies in both Old and New World primates (Boussaoud et al., 1992; Palmer and Rosa, 2006; Kaas and Lyon, 2007).

In contrast, projections from inferior temporal areas TE3, TE2, and TEO were generally sparse. Except for subject M1551 (**Fig. 3J, K**), retrograde labelling in these regions was limited, with modest labelling observed in subjects F1746 (**Fig. 2J, K**) and M1923 (**Fig. 3.5J, K**), and near absence of label in M1842 (**Fig. 4J–L**). Although the distribution of projections within inferior temporal cortex varied across subjects, previous studies involving broad tracer injections in TE and TEO in the macaque have demonstrated projections from inferior temporal cortex to PIm (Baleydier and Morel, 1992; Baizer et al., 1993; Webster et al., 1993).

**Figure 3:**
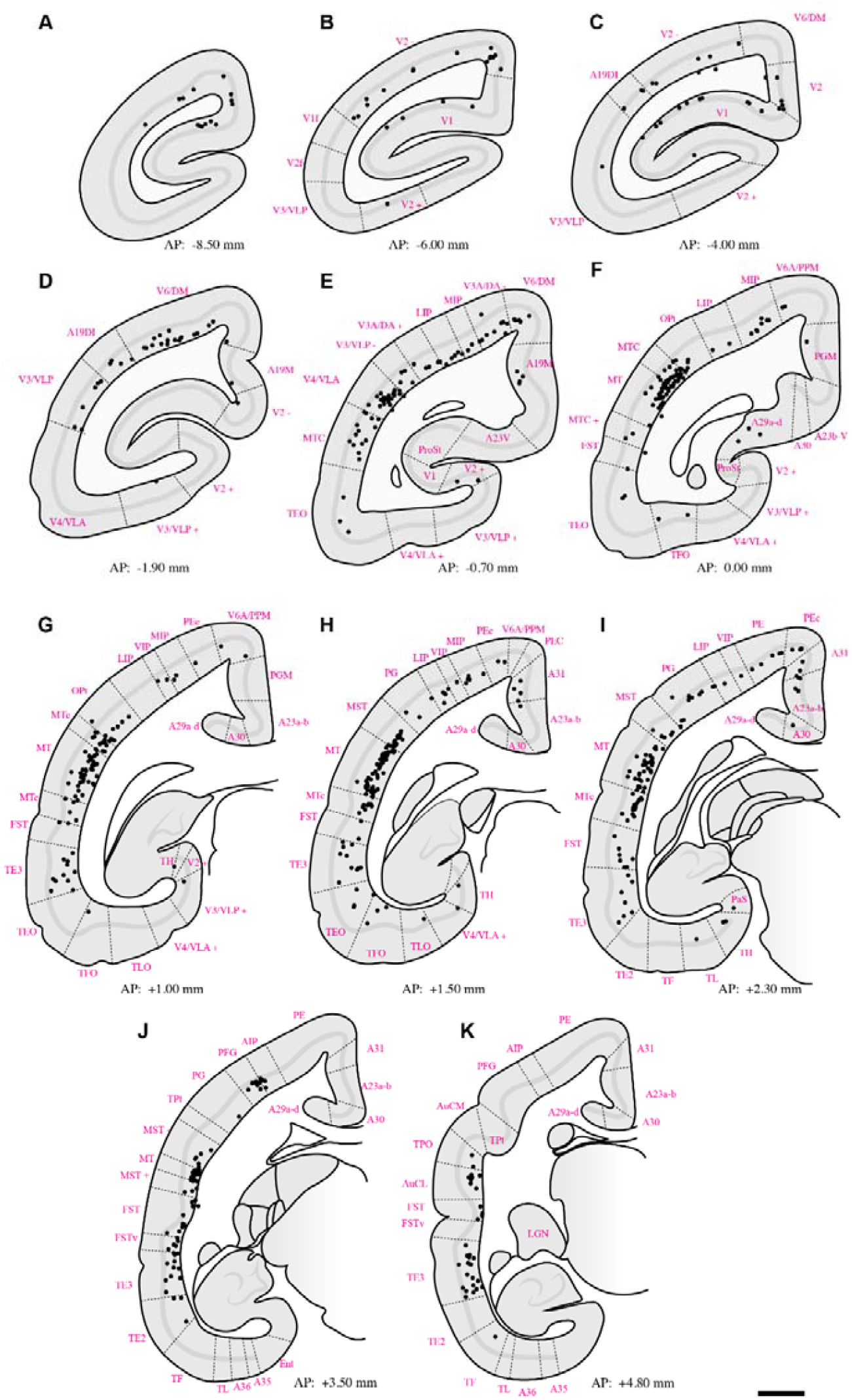
Summary of retrograde label throughout case **M1551** following deposition of tracer in PIm (refer to Fig 1G for tracer bolus placement). Dotted lines indicate boundaries of cortical areas. Abbreviations of cortical areas summarised in Table 1. Scale bar = 2mm.

**Figure 4:**
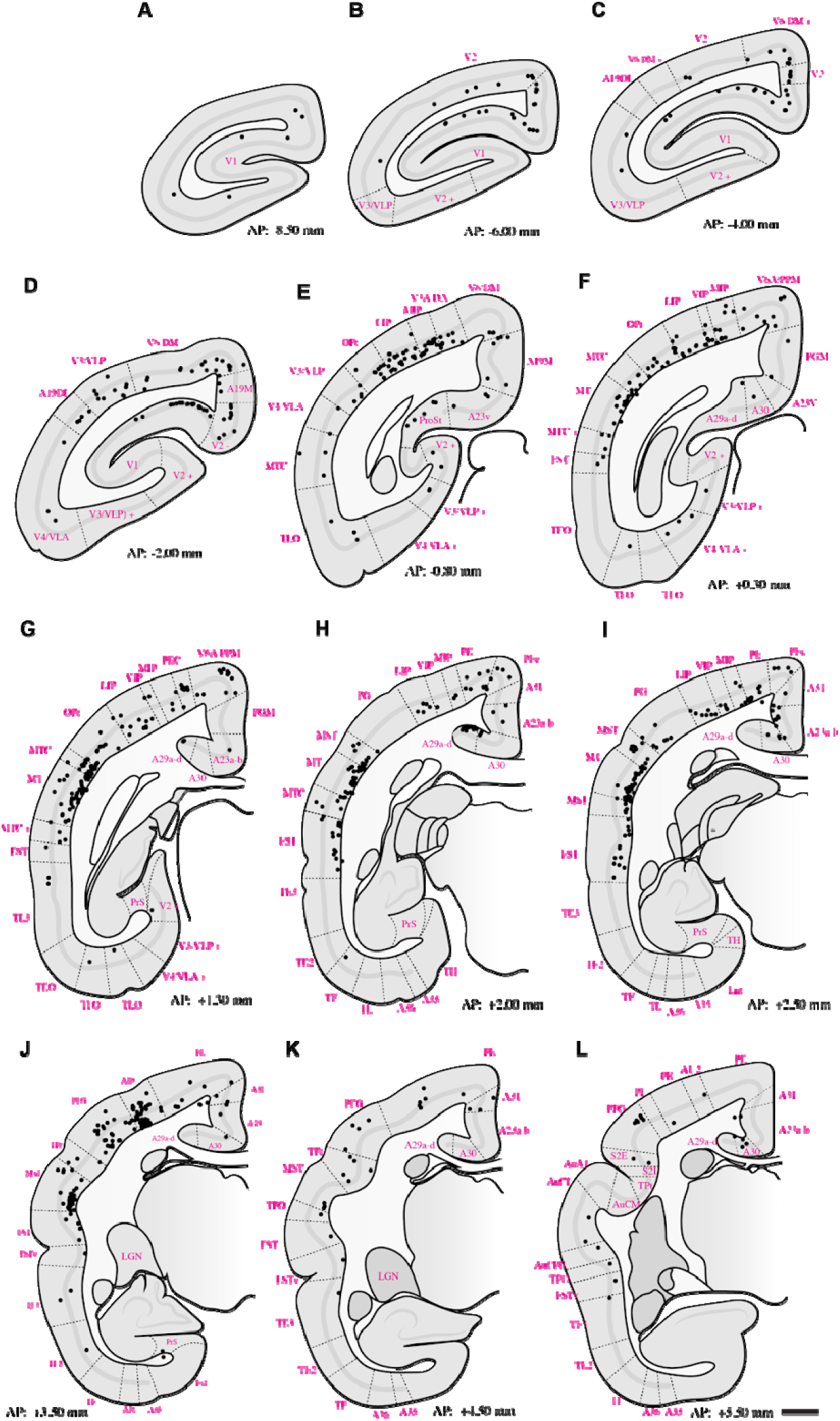
Summary of retrograde label throughout case **M1842** following deposition of tracer in PIm (refer to Fig 1G for tracer bolus placement). Dotted lines indicate boundaries of cortical areas. Abbreviations of cortical areas summarised in Table 1. Scale bar = 2mm.

**Figure 5:**
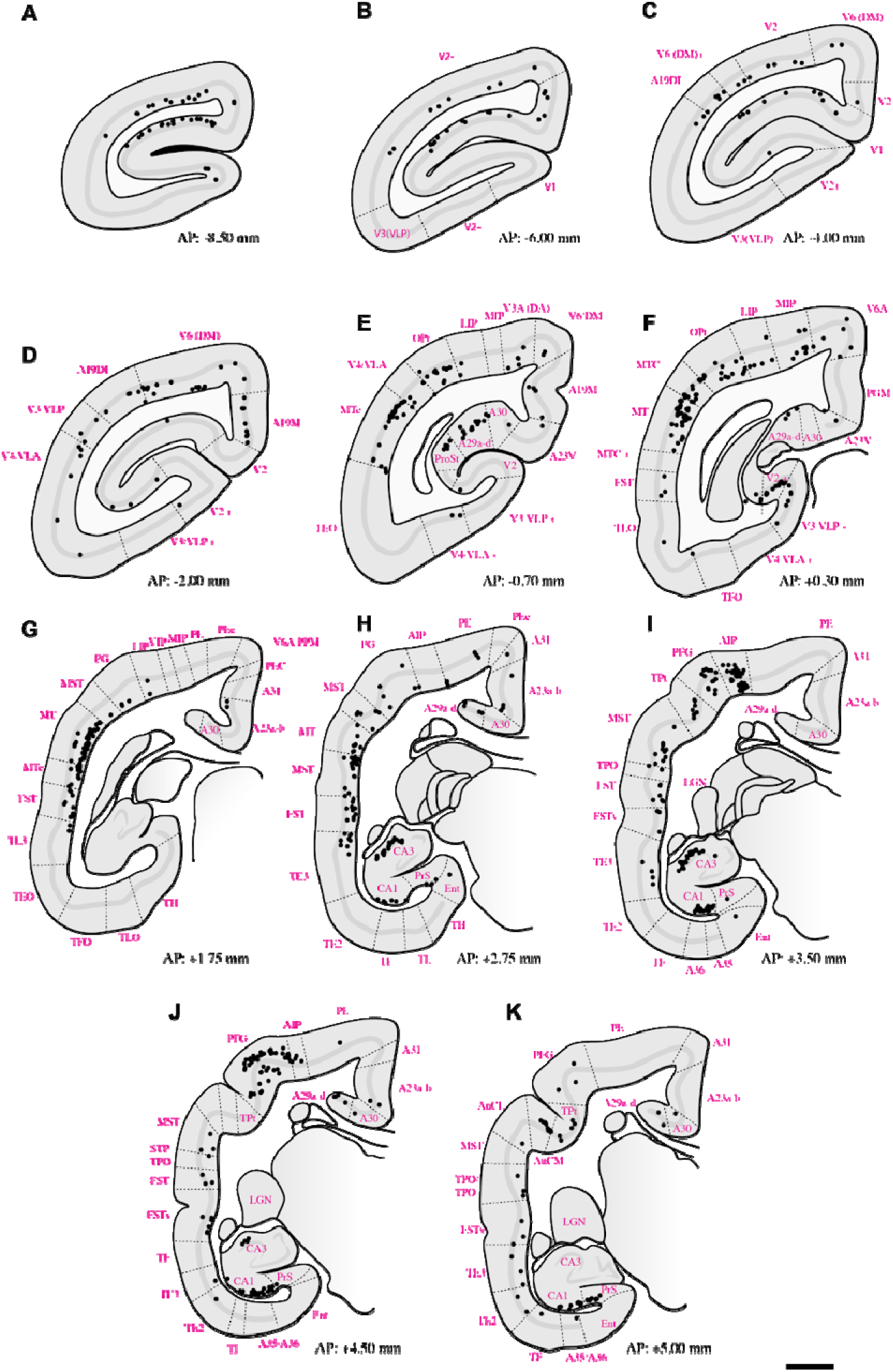
Summary of retrograde label throughout case **M1923** following deposition of tracer in PIm (refer to Fig 1G for tracer bolus placement). Dotted lines indicate boundaries of cortical areas. Abbreviations of cortical areas summarised in Table 1. Scale bar = 2mm.

Within the parietal cortex, retrograde labelling was observed in regions surrounding the intraparietal sulcus, including AIP, MIP, LIP, and VIP, as well as in superior parietal areas PE and PEc, inferior parietal areas PF, PFG, and PG, and the occipitoparietal transitional area (OPt). Across subjects, the distribution of PIm-projecting neurons in MIP, LIP, and VIP was consistent, forming a mediolaterally distributed band of retrogradely labelled cells (**Fig. 2–5: E– G**). In three of four cases (**Fig. 2I; Fig. 4J; Fig. 5I**), AIP exhibited comparatively denser retrograde labelling than other intraparietal areas.

The presence of direct projections from LIP to PIm is consistent with findings in the macaque (Giolli et al., 2001). To our knowledge, this is the first report demonstrating connectivity between PIm and VIP, MIP, and AIP in the marmoset. Previous tract-tracing studies in Old World macaques have generally reported projections from intraparietal regions to the anterior or medial pulvinar nuclei (Baizer et al., 1993; Padberg and Krubitzer, 2006; Cappe et al., 2007; Prevosto et al., 2009).

Projections to PIm from inferior parietal areas were generally more extensive than those from superior parietal regions across all four cases (**Fig. 2–5E–K**). Within the inferior parietal cortex, retrograde labelling was consistently denser in OPt and PFG compared to PF and PG. Across subjects, the MT complex—comprising MT, MTc, and MST—typically exhibited the densest retrograde labelling overall (**Fig. 2–5F–H**). An exception was observed in case F1746, in which more rostral representations within the superior posterior parietal cortex (V6A/PPM, PEc, and PE) also showed substantial projections to PIm (**Fig. 2I,J**). Sparse retrograde labelling in PEc was observed in three of four cases (**Fig. 2G; Fig. 3–4I**).

Previous studies examining posterior parietal projections to the pulvinar have reported variable findings. Several investigations have identified the medial pulvinar (PM) as the principal recipient of posterior parietal inputs, with minimal contribution from PI subdivisions (Yeterian and Pandya, 1985; Baleydier and Mauguière, 1987; Baleydier and Morel, 1992; Impieri et al., 2018). In contrast, other studies have reported projections from parietal cortex to PIm (Kasdon and Jacobson, 1978; Brysch et al., 1990; Baizer et al., 1993).

Within the posterior parietal cortex (PPC), retrograde labelling was not uniformly distributed, suggesting that projections to PIm may arise from specific topographic subdivisions. A recent tract-tracing study in the prosimian galago reported that projections to PI originate predominantly from the dorsal-caudal PPC (Wang et al., 2023), a region implicated in visually-guided reaching behaviours. Notably, this subdivision receives input from peripheral representations of extrastriate visual cortex (Stepniewska et al., 2015). In humans, parietal lesions have been associated with impairments in reaching toward targets in the peripheral visual field, with deficits reduced following foveation (Passarelli et al., 2021). Together, these findings provide a comparative framework for interpreting the pattern of parietal projections to PIm observed in the present study.

### 3.4 Projections from other domains

Across all cases, retrograde labelling was observed in the retrosplenial cortex (areas A29 and A30) and posterior cingulate regions (A23a–b, A31, and PGM). In addition, retrogradely labelled neurons were identified in hippocampal subfields CA1 and CA3 in two cases (F1746 and M1923) which had tracer spread extended into ventral portions of PM.

Previous tract-tracing studies have primarily reported connectivity between retrosplenial, posterior cingulate cortex or hippocampus with the medial (PM) or lateral (PL) subdivisions of the pulvinar, with no clear evidence of projections to PI subdivisions (Baleydier and Mauguiere, 1985; Leichnetz, 2001; Shibata and Yukie, 2003; Buckwalter et al., 2008; Gamberini et al., 2020; Togo et al., 2025). The densest retrosplenial labelling was observed in subjects F1746 (**Fig. 2F– I**) and M1923 (**Fig. 5E,F**), the cases tracer spread into medial pulvinar. However, in case M1842, where tracer deposition was confined to PIm, whilst CA1 and CA3 label was absent, substantial retrograde labelling was also present in posterior cingulate and retrosplenial cortices (**Fig. 4H,I**).

### 3.5 Semi-quantitative assessment of cortical connectivity with PIm

To assess the relative strength of cortical inputs to PIm, we quantified the number of retrogradely labelled neurons in each cortical area as a proportion of the total labelled cells observed per subject (**Fig. 6**). Consistent with previous reports indicating that PI subdivisions are predominantly associated with visual processing, inputs from early visual areas in the occipital lobe and the MT complex in the temporal lobe accounted for nearly 60% of total retrograde labelling. V1 and MT which act as the primary early maturing nodes of the visual cortex (Bourne, 2010) together contributed approximately 25% of all cortical inputs to PIm.

**Figure 6:**
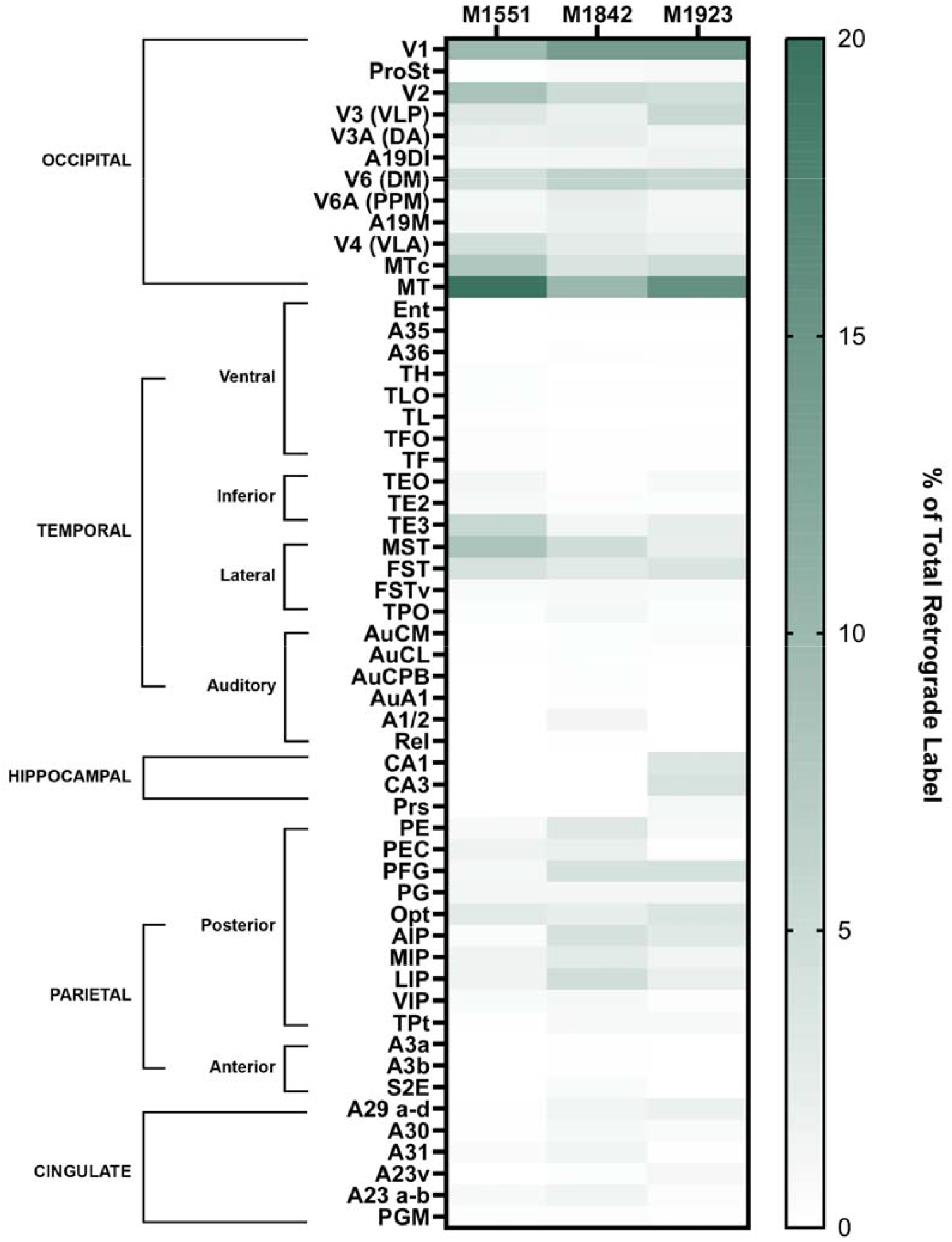
Semi-quantitative assessment of cortical contributions to PIm, calculated as the number of labeled cells in each area divided by the total number of labeled cells quantified within each subject. Areas are grouped by major regions and subregions according to the marmoset brain atlas (Paxinos et al., 2012; Córdoba-Claros et al., 2025). See Table 1 for area abbreviations.

The parietal cortex represented the second largest source of cortical input, contributing close to 20% of total retrograde label. In contrast to the occipital domain, where V1 and MT showed markedly greater labelling than other visual areas, the distribution of labelled neurons within the parietal cortex was comparatively uniform. Areas PFG, OPt, and LIP each accounted for approximately 3% of total labelled cells.

Retrograde labelling was also consistently observed within cingulate regions across subjects, contributing approximately 4% of the total cortical input. Contributions from the temporal cortex were more selective; while most inferior temporal areas showed limited labelling, TE3, MST, and FST together comprised over 10% of total cortical projections to PIm. In two of four cases, retrogradely labelled neurons were additionally identified within the hippocampal formation.

## 4 DISCUSSION

Our goal in undertaking this study was to reconcile a fragmented literature and move the field toward a clearer consensus on corticopulvinar circuits centered on the medial subdivision of the inferior pulvinar (PIm). Much of what we currently understand about cortical–pulvinar connectivity derives from studies in Old World monkeys, where key cortical territories are buried deep within sulci and can be challenging to access comprehensively. As the pattern of cortical inputs to a thalamic nucleus provides critical clues to its functional identity, a systematic and targeted mapping of cortical projections to PIm was needed. By placing retrograde tracers directly within PIm, we aimed to generate a more complete cortical–PIm connectivity map and, in doing so, reveal functional contributions that may have remained underappreciated

Our retrograde tracer injections into the peripheral PIm identified a distributed constellation of cortical areas projecting to this thalamic region. Considering the mosaic organization of primate visual cortex, where distinct areas exhibit eccentricity biases, some preferentially representing the fovea (Rosa and Tweedale, 2000), while others emphasize peripheral visual space (Palmer and Rosa, 2006), these findings help clarify which cortical domains preferentially engage the peripheral sector of PIm. This distinction is particularly relevant given the disproportionate role of foveal processing in primate vision. An important unresolved question is whether the foveal portion of PIm (the ‘calbindin-negative tail’) shares a similar connectional architecture or instead participates in a distinct corticopulvinar network. Prior work suggests that direct input from foveal representations of early visual cortex may be sparse, as tracer injections restricted to the marmoset V1 operculum (0–5° eccentricity) did not label PIm (Kaas and Lyon, 2007). However, anterograde tracer placements near the V1–V2 border have revealed axonal terminations within PIm (Gutierrez and Cusick, 1997), leaving open the possibility that specialized corticopulvinar circuits selectively link foveal cortex with this thalamic territory.

Several limitations should be noted. The first is that only one case had tracer restricted entirely in PIm. Most notably, in two cases, some tracer encroached on the ventral portion of PM. In these two cases alone, we observed projections from the hippocampus, and in the case where PIcm was also injected, this subject had a higher proportion of cortical inputs from the inferior temporal domain. Nevertheless, after normalizing the total retrograde label in each subject, that it was clear the distribution of cortical afferents across all subjects converged to show a similar spatial profile **(Fig 6)**. The other limitation worth highlighting is that the extensive cortical labeling observed after PIm injections precluded a fully reciprocal strategy in which tracers would be placed in every retrogradely labeled cortical area. Instead, we interpreted our findings in the context of the substantial corticopulvinar literature in monkeys, where tracers were typically administered in defined cortical fields. Our analysis also prioritized occipital and visual cortices; technical limitations during cryosectioning limited recovery of the frontal pole, and thus, potential rostral cortical inputs to PIm were not systematically examined. Frontal projections to the pulvinar are generally reported to converge within the medial pulvinar (Brysch et al., 1990; Romanski et al., 1997; Roberts et al., 2007; Homman-Ludiye et al., 2019; Córdoba-Claros et al., 2025), although sparse projections from PI and PIm to areas 8A and 46v have been described (Romanski et al., 1997; Córdoba-Claros et al., 2025). The opportunity to reveal additional frontal inputs to PIm appears unlikely, given the visual specialization of PI, although a truly comprehensive account of corticopulvinar organization will ultimately require direct examination of rostral cortices. Species differences also merit consideration. Most pulvinocortical tracing studies have been conducted in macaques, and discrepancies with prior reports (e.g., parietal or retrosplenial connectivity with PIm) may reflect marmoset-specific organization. Whilst species-specific attributes will always be inevitable, methods such as consensus mapping, a technique that unifies species-specific anatomical atlases using a common parcellation scheme, offer an elegant solution for directly interrogating commonalities and differences between primate species (Magrou et al., 2026). Supporting this possibility, functional imaging in marmosets demonstrated activation of PI and PM subdivisions, along with occipital, parietal, temporal, inferior temporal, and retrosplenial cortices, during looming stimulation (Cléry et al., 2020), although fMRI cannot resolve specific corticopulvinar circuits at the PI subdivision level. As the marmoset model gains prominence for its lissencephalic cortex (Solomon and Rosa, 2014) and its suitability for social neuroscience (Miller et al., 2016; Scott and Bourne, 2022), accurate species-specific connectional maps are increasingly important. Future tracer studies targeting underexplored regions, including medial extrastriate and posterior cingulate cortices, will be necessary to refine the corticopulvinar framework proposed here.

Collectively, our findings position PIm as a central integrative node within the primate visual system, receiving convergent input from occipital, temporal, and parietal cortices. This extensive connectivity reinforces the emerging view of the pulvinar as an active regulator of cortical communication, dynamically shaping excitatory and inhibitory interactions across distributed networks (Kohn et al., 2020). The distribution of retrograde labeling indicates that PIm incorporates signals from multiple tiers of the visual hierarchy, including the middle temporal complex, a core substrate for motion processing (Rosa and Elston, 1998). Importantly, projections from intraparietal regions involved in reach and grasp (Vesia and Crawford, 2012), superior posterior parietal areas supporting visuomotor guidance and body-in-space awareness (Passarelli et al., 2021), and inferior parietal territories implicated in extrapersonal spatial processing (Kravitz et al., 2011) extend PIm’s function beyond perception and into action-oriented visual domains.

Viewed in this light, PIm is not merely a relay within the visual thalamus but a nexus where perceptual, spatial, and visuomotor signals converge, but a flexible and plastic node that could enable the coordination of sensory inputs with behavioral demands. Although retrosplenial and posterior cingulate cortices, which contribute to navigation and spatial memory (Vann et al., 2009; Leech and Smallwood, 2019), remain less clearly linked to PIm in our data, an even broader sensory integration repertoire remains a possibility. Taken together, the connectional architecture described here supports a model in which PIm participates in orchestrating multidimensional visual representations, aligning motion, space, and action within a unified thalamocortical framework.

Existing circuit-level evidence further supports an integrative role for PIm. This subdivision receives input from diverse classes of wide-field retinal ganglion cells that are sensitive to visual motion, looming stimuli, and signals relevant for visual pursuit (Puller et al., 2015; Grünert et al., 2021; Kim et al., 2022), positioning it to access rapid, behaviorally salient visual information. PIm has also been implicated in the development of goal-directed reach and grasp behaviours (Mundinano et al., 2018) and in filtering visual distractors (Snow et al., 2009), highlighting its contribution to action selection and attentional control. Considered alongside the extensive cortical inputs identified here, from motion-sensitive visual areas and parietal visuomotor regions, PIm appears strategically situated as the conduit for perception and action. Rather than functioning as a passive relay, PIm may serve as an early thalamic gatekeeper that primes and stabilizes visuomotor networks, aligning salient visual signals with the coordinated execution of hand–eye behaviors such as reach and grasp (**Fig. 7**).

**Figure 7:**
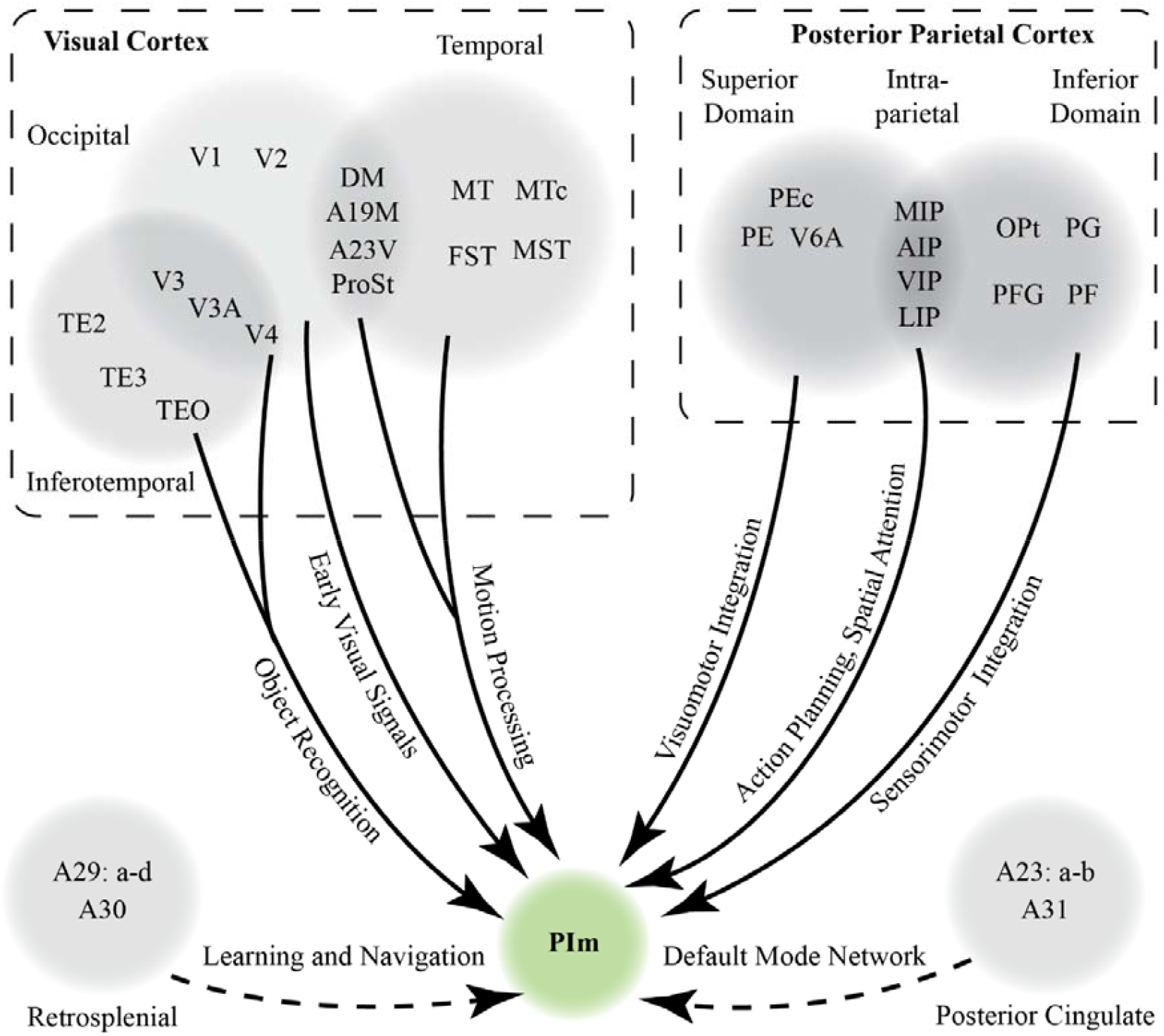
Summary of the retrograde tracing results in this study and our proposed network of PIm cortical connectivity. Solid lines denotes the functional domains PIm receives input from which were observed in this study and supported by other previous literature in the nonhuman primate. Dashed lines represent projections from areas which are not definitive due to tracer encroaching in adjacent areas and the lack of previous literature reporting afferents from that domain.

Taken together, our findings support a model in which the inferior pulvinar, particularly the PIm subdivision, serves as a principal hub for coordinating corticocortical interactions. The cortico-thalamo-cortical loops enable the observer to rapidly interpret and anticipate appropriate motor responses to dynamic visual input, consistent with the action-oriented functions ascribed to the dorsal stream (Goodale and Milner, 1992). Beyond visuomotor integration, connectivity between PI and the amygdala, a pathway implicated in processing fearful and behaviorally salient stimuli (McFadyen et al., 2019) raises the possibility that the inferior pulvinar operates as an early responder to biologically relevant sensory events (Soares et al., 2017). In natural environments, visual stimuli vary widely in emotional and behavioral significance, often demanding responses that extend beyond stereotyped reach-and-grasp actions to include vocalization, defensive reactions, or higher-order cognitive strategies. Although PI lacks strong direct projections to frontal cortex, its influence may nevertheless propagate to executive networks through short polysynaptic routes. Notably, the PM nucleus projects robustly to frontal cognitive centers, and overlapping temporal (e.g., TE3, TE2) and parietal (e.g., PFG, PG) territories interface with both PI and PM. These shared circuit motifs suggest that PI and PM cooperate via distributed temporal–parietal loops to regulate information flow between early visual areas and higher-order cognitive systems. Such a framework resonates with the global neuronal workspace model (Cortes et al., 2024), in whichu ascending pulvinar projections amplify and sustain cortical signals, preventing their attenuation before they reach distributed cognitive networks. Testing these hypotheses will require technically demanding circuit-level manipulations, yet emerging genetic technologies, coupled with increasingly sophisticated quantification of naturalistic primate behavior, now make such investigations feasible. By establishing a comprehensive anatomical framework of the PIm network, the present study lays critical groundwork for future efforts to decipher how the pulvinar coordinates brain-wide communication, transforming transient sensory signals into coherent, adaptive behavior.

## Acknowledgements

The authors would like to acknowledge and thank the Monash Micro Imaging Platform for providing microscopy resources and technical assistance. This research was supported, in part, by the Intramural Research Program of the National Institutes of Health (NIH). The contributions of the NIH author(s) are considered Works of the United States Government. The findings and conclusions presented in this paper are those of the author(s) and do not necessarily reflect the views of the NIH or the U.S. Department of Health and Human Services.

## REFERENCES

Arafune-Mishima A, Abe H, Tani T, Mashiko H, Watanabe S, Sakai K, Suzuki W, Mizukami H, Watakabe A, Yamamori T, Ichinohe N (2020) Axonal Projections from Middle Temporal Area to the Pulvinar in the Common Marmoset. Neuroscience 446:145–156.

Asanuma C, Andersen RA, Cowan WM (1985) The thalamic relations of the caudal inferior parietal lobule and the lateral prefrontal cortex in monkeys: Divergent cortical projections from cell clusters in the medial pulvinar nucleus. Journal of Comparative Neurology 241:357–381.

Baizer JS, Desimone R, Ungerleider LG (1993) Comparison of subcortical connections of inferior temporal and posterior parietal cortex in monkeys. Visual Neuroscience 10:59–72.

Baldwin MKL, Bourne JA (2020) Chapter 24 - The Evolution of Subcortical Pathways to the Extrastriate Cortex. In: Evolutionary Neuroscience (Second Edition) (Kaas JH, ed), pp 565–587. London: Academic Press.

Baldwin MKL, Kaskan PM, Zhang B, Chino YM, Kaas JH (2012) Cortical and subcortical connections of V1 and V2 in early postnatal macaque monkeys. Journal of Comparative Neurology 520:544–569.

Baleydier C, Mauguiere F (1985) Anatomical evidence for medial pulvinar connections with the posterior cingulate cortex, the retrosplenial area, and the posterior parahippocampal gyrus in monkeys. Journal of Comparative Neurology 232:219–228.

Baleydier C, Mauguière F (1987) Network organization of the connectivity between parietal area 7, posterior cingulate cortex and medial pulvinar nucleus: a double fluorescent tracer study in monkey. Experimental Brain Research 66:385–393.

Baleydier C, Morel A (1992) Segregated thalamocortical pathways to inferior parietal and inferotemporal cortex in macaque monkey. Visual Neuroscience 8:391–405.

Benevento LA, Rezak M (1976) The cortical projections of the inferior pulvinar and adjacent lateral pulvinar in the rhesus monkey (macaca mulatta): An autoradiographic study. Brain Research 108:1–24.

Born RT, Bradley DC (2005) Strucuture and Function of Visual Area MT. Annual Review of Neuroscience 28:157–189.

Bourne JA (2010) Unravelling the development of the visual cortex: implications for plasticity and repair. Journal of Anatomy 217:449–468.

Bourne JA, Rosa MGP (2005) Hierarchical Development of the Primate Visual Cortex, as Revealed by Neurofilament Immunoreactivity: Early Maturation of the Middle Temporal Area (MT). Cerebral Cortex 16:405–414.

Boussaoud D, Desimone R, Ungerleider LG (1992) Subcortical connections of visual areas MST and FST in macaques. Visual Neuroscience 9:291–302.

Brysch W, Brysch I, Creutzfeldt OD, Schlingensiepen R, Schlingensiepen KH (1990) The topology of the thalamo-cortical projections in the marmoset monkey (Callithrix jacchus). Experimental Brain Research 81:1–17.

Buckwalter JA, Parvizi J, Morecraft RJ, van Hoesen GW (2008) Thalamic projections to the posteromedial cortex in the macaque. Journal of Comparative Neurology 507:1709–1733.

Cappe C, Morel A, Rouiller EM (2007) Thalamocortical and the dual pattern of corticothalamic projections of the posterior parietal cortex in macaque monkeys. Neuroscience 146:1371–1387.

Cléry JC, Schaeffer DJ, Hori Y, Gilbert KM, Hayrynen LK, Gati JS, Menon RS, Everling S (2020) Looming and receding visual networks in awake marmosets investigated with fMRI. NeuroImage 215:116815.

Córdoba-Claros MA, Rubio-Garrido P, de Lima RRM, Morais PLAG, do Nascimento ES, Cavalcante JS, Clascá F (2025) Projection Motifs and Wiring Logic of Medial Pulvinar Thalamocortical Axons in the Marmoset Monkey. The Journal of Neuroscience 45:e1837242025.

Cortes N, Ladret HJ, Abbas-Farishta R, Casanova C (2024) The pulvinar as a hub of visual processing and cortical integration. Trends in Neurosciences 47:120–134.

Cusick CG, Scripter JL, Darensbourg JG, Weber JT (1993) Chemoarchitectonic subdivisions of the visual pulvinar in monkeys and their connectional relations with the middle temporal and rostral dorsolateral visual areas, MT and DLr. Journal of Comparative Neurology 336:1–30.

Diamond IT, Fitzpatrick D, Conley M (1992) A projection from the parabigeminal nucleus to the pulvinar nucleus in Galago. Journal of Comparative Neurology 316:375–382.

Dick A, Kaske A, Creutzfeldt OD (1991) Topographical and topological organization of the thalamocortical projection to the striate and prestriate cortex in the marmoset (Callithrix jacchus). Experimental Brain Research 84:233–253.

Felleman DJ, Van Essen DC (1991) Distributed Hierarchical Processing in the Primate Cerebral Cortex. Cerebral Cortex 1:1–47.

Fitzgibbon T, Szmajda BA, Martin PR (2007) First order connections of the visual sector of the thalamic reticular nucleus in marmoset monkeys (Callithrix jacchus). Visual Neuroscience 24:857–874.

Fox DM, Goodale MA, Bourne JA (2020) The Age-Dependent Neural Substrates of Blindsight. Trends in Neurosciences 43:242–252.

Fritsches KA, Rosa MGP (1996) Visuotopic organisation of striate cortex in the marmoset monkey (Callithrix jacchus). Journal of Comparative Neurology 372:264–282.

Gamberini M, Passarelli L, Impieri D, Worthy KH, Burman KJ, Fattori P, Galletti C, Rosa MGP, Bakola S (2020) Thalamic afferents emphasize the different functions of macaque precuneate areas. Brain Structure and Function 225:853–870.

Gattass R, Soares JGM, Lima B (2018) Connectivity of the Pulvinar. In: The Pulvinar Thalamic Nucleus of Non-Human Primates: Architectonic and Functional Subdivisions, pp 19–29. Cham: Springer International Publishing.

Gattass R, Galkin TW, Desimone R, Ungerleider LG (2014) Subcortical connections of area V4 in the macaque. Journal of Comparative Neurology 522:1941–1965.

Giolli RA, Gregory KM, Suzuki DA, Blanks RH, Lui F, Betelak KF (2001) Cortical and subcortical afferents to the nucleus reticularis tegmenti pontis and basal pontine nuclei in the macaque monkey. Vis Neurosci 18:725–740.

Goodale MA, Milner AD (1992) Separate visual pathways for perception and action. Trends in Neurosciences 15:20–25.

Grünert U, Lee SCS, Kwan WC, Mundinano I-C, Bourne JA, Martin PR (2021) Retinal ganglion cells projecting to superior colliculus and pulvinar in marmoset. Brain Structure and Function 226:2745–2762.

Guedj C, Vuilleumier P (2023) Modulation of pulvinar connectivity with cortical areas in the control of selective visual attention. NeuroImage 266:119832.

Gutierrez C, Cusick CG (1997) Area V1 in macaque monkeys projects to multiple histochemically defined subdivisions of the inferior pulvinar complex. Brain Research 765:349–356.

Homman-Ludiye J, Mundinano IC, Kwan WC, Bourne JA (2019) Extensive Connectivity Between the Medial Pulvinar and the Cortex Revealed in the Marmoset Monkey. Cerebral Cortex 30:1797–1812.

Huo B-X, Zeater N, Lin MK, Takahashi YS, Hanada M, Nagashima J, Lee BC, Hata J, Zaheer A, Grünert U, Miller MI, Rosa MGP, Okano H, Martin PR, Mitra PP (2019) Relation of koniocellular layers of dorsal lateral geniculate to inferior pulvinar nuclei in common marmosets. European Journal of Neuroscience 50:4004–4017.

Impieri D, Gamberini M, Passarelli L, Rosa MGP, Galletti C (2018) Thalamo-cortical projections to the macaque superior parietal lobule areas PEc and PE. Journal of Comparative Neurology 526:1041–1056.

Kaas JH, Lyon DC (2007) Pulvinar contributions to the dorsal and ventral streams of visual processing in primates. Brain Research Reviews 55:285–296.

Kasdon DL, Jacobson S (1978) The thalamic afferents to the inferior parietal lobule of the rhesus monkey. Journal of Comparative Neurology 177:685–705.

Kennedy H, Bullier J (1985) A double-labeling investigation of the afferent connectivity to cortical areas V1 and V2 of the macaque monkey. The Journal of Neuroscience 5:2815–2830.

Kim YJ, Peterson BB, Crook JD, Joo HR, Wu J, Puller C, Robinson FR, Gamlin PD, Yau K-W, Viana F, Troy JB, Smith RG, Packer OS, Detwiler PB, Dacey DM (2022) Origins of direction selectivity in the primate retina. Nature Communications 13:2862.

Kinoshita M, Kato R, Isa K, Kobayashi K, Kobayashi K, Onoe H, Isa T (2019) Dissecting the circuit for blindsight to reveal the critical role of pulvinar and superior colliculus. Nature Communications 10:135.

Kohn A, Jasper AI, Semedo JD, Gokcen E, Machens CK, Yu BM (2020) Principles of Corticocortical Communication: Proposed Schemes and Design Considerations. Trends in Neurosciences 43:725–737.

Kosslyn SM, Thompson WL (2003) When is early visual cortex activated during visual mental imagery? Psychological Bulletin 129:723–746.

Kravitz DJ, Saleem KS, Baker CI, Mishkin M (2011) A new neural framework for visuospatial processing. Nature Reviews Neuroscience 12:217–230.

Kwan WC, Mundinano I-C, de Souza MJ, Lee SCS, Martin PR, Grünert U, Bourne JA (2019) Unravelling the subcortical and retinal circuitry of the primate inferior pulvinar. Journal of Comparative Neurology 527:558–576.

Leech R, Smallwood J (2019) Chapter 5 - The posterior cingulate cortex: Insights from structure and function. In: Handbook of Clinical Neurology (Vogt BA, ed), pp 73-85: Elsevier.

Leichnetz GR (2001) Connections of the medial posterior parietal cortex (area 7m) in the monkey. The Anatomical Record 263:215–236.

Magrou L, Theodoni P, Arnsten AFT, Rosa MGP, Wang X-J (2026) From comparative connectomics to large-scale working memory modeling in macaque and marmoset. Cell Reports 45:116847.

Marion R, Li K, Purushothaman G, Jiang Y, Casagrande VA (2013) Morphological and neurochemical comparisons between pulvinar and V1 projections to V2. Journal of Comparative Neurology 521:813–832.

McFadyen J, Mattingley JB, Garrido MI (2019) An afferent white matter pathway from the pulvinar to the amygdala facilitates fear recognition. eLife 8:e40766.

Mikellidou K, Kurzawski JW, Frijia F, Montanaro D, Greco V, Burr DC, Morrone MC (2017) Area Prostriata in the Human Brain. Current Biology 27:3056-3060.e3053.

Miller Cory T, Freiwald Winrich A, Leopold David A, Mitchell Jude F, Silva Afonso C, Wang X (2016) Marmosets: A Neuroscientific Model of Human Social Behavior. Neuron 90:219–233.

Mundinano I-C, Flecknell PA, Bourne JA (2016) MRI-guided stereotaxic brain surgery in the infant and adult common marmoset. Nature Protocols 11:1299–1308.

Mundinano I-C, Kwan WC, Bourne JA (2019) Retinotopic specializations of cortical and thalamic inputs to area MT. Proceedings of the National Academy of Sciences 116:23326–23331.

Mundinano I-C, Fox DM, Kwan WC, Vidaurre D, Teo L, Homman-Ludiye J, Goodale MA, Leopold DA, Bourne JA (2018) Transient visual pathway critical for normal development of primate grasping behavior. Proceedings of the National Academy of Sciences 115:1364–1369.

Münkle MC, Waldvogel HJ, Faull RLM (2000) The distribution of calbindin, calretinin and parvalbumin immunoreactivity in the human thalamus. Journal of Chemical Neuroanatomy 19:155–173.

O’Brien BJ, Abel PL, Olavarria JF (2001) The retinal input to calbindin-D28k-defined subdivisions in macaque inferior pulvinar. Neuroscience Letters 312:145–148.

Ogren MP, Hendrickson AE (1977) The distribution of pulvinar terminals in visual areas 17 and 18 of the monkey. Brain Research 137:343–350.

Ohye C (2002) Thalamus and Thalamic Damage. In: Encyclopedia of the Human Brain (Ramachandran VS, ed), pp 575–597. New York: Academic Press.

Padberg J, Krubitzer L (2006) Thalamocortical connections of anterior and posterior parietal cortical areas in New World titi monkeys. Journal of Comparative Neurology 497:416–435.

Palmer SM, Rosa MGP (2006) A distinct anatomical network of cortical areas for analysis of motion in far peripheral vision. European Journal of Neuroscience 24:2389–2405.

Passarelli L, Gamberini M, Fattori P (2021) The superior parietal lobule of primates: a sensory-motor hub for interaction with the environment. J Integr Neurosci 20:157–171.

Paxinos G, Watson C, Petrides M, Rosa M, Tokuno H (2012) The marmoset brain in stereotaxic coordinates, 1st Edition. London ;: Academic Press.

Prevosto V, Graf W, Ugolini G (2009) Posterior parietal cortex areas MIP and LIPv receive eye position and velocity inputs via ascending preposito-thalamo-cortical pathways. European Journal of Neuroscience 30:1151–1161.

Puller C, Manookin MB, Neitz J, Rieke F, Neitz M (2015) Broad Thorny Ganglion Cells: A Candidate for Visual Pursuit Error Signaling in the Primate Retina. The Journal of Neuroscience 35:5397–5408.

Roberts AC, Tomic DL, Parkinson CH, Roeling TA, Cutter DJ, Robbins TW, Everitt BJ (2007) Forebrain connectivity of the prefrontal cortex in the marmoset monkey (Callithrix jacchus): An anterograde and retrograde tract-tracing study. Journal of Comparative Neurology 502:86–112.

Romanski LM, Giguere M, Bates JF, Goldman-Rakic PS (1997) Topographic organization of medial pulvinar connections with the prefrontal cortex in the rhesus monkey. Journal of Comparative Neurology 379:313–332.

Rosa MGP, Elston GN (1998) Visuotopic organisation and neuronal response selectivity for direction of motion in visual areas of the caudal temporal lobe of the marmoset monkey (Callithrix jacchus): Middle temporal area, middle temporal crescent, and surrounding cortex. Journal of Comparative Neurology 393:505–527.

Rosa MGP, Tweedale R (2000) Visual areas in lateral and ventral extrastriate cortices of the marmoset monkey. Journal of Comparative Neurology 422:621–651.

Saalmann YB, Pinsk MA, Wang L, Li X, Kastner S (2012) The Pulvinar Regulates Information Transmission Between Cortical Areas Based on Attention Demands. Science 337:753–756.

Schindelin J, Arganda-Carreras I, Frise E, Kaynig V, Longair M, Pietzsch T, Preibisch S, Rueden C, Saalfeld S, Schmid B, Tinevez J-Y, White DJ, Hartenstein V, Eliceiri K, Tomancak P, Cardona A (2012) Fiji: an open-source platform for biological-image analysis. Nature Methods 9:676–682.

Scott JT, Bourne JA (2022) Modelling behaviors relevant to brain disorders in the nonhuman primate: Are we there yet? Progress in Neurobiology 208:102183.

Sherman SM, Guillery RW (1998) On the actions that one nerve cell can have on another: Distinguishing “drivers” from “modulators”. Proceedings of the National Academy of Sciences 95:7121–7126.

Sherman SM, Guillery RW (2002) The role of the thalamus in the flow of information to the cortex. Philosophical Transactions of the Royal Society B: Biological Sciences 357:1695–1708.

Shibata H, Yukie M (2003) Differential thalamic connections of the posteroventral and dorsal posterior cingulate gyrus in the monkey. European Journal of Neuroscience 18:1615–1626.

Snow JC, Allen HA, Rafal RD, Humphreys GW (2009) Impaired attentional selection following lesions to human pulvinar: Evidence for homology between human and monkey. Proceedings of the National Academy of Sciences 106:4054–4059.

Soares SC, Maior RS, Isbell LA, Tomaz C, Nishijo H (2017) Fast Detector/First Responder: Interactions between the Superior Colliculus-Pulvinar Pathway and Stimuli Relevant to Primates. Frontiers in Neuroscience Volume 11 -2017.

Solomon SG, Rosa MGP (2014) A simpler primate brain: the visual system of the marmoset monkey. Frontiers in Neural Circuits Volume 8 -2014.

Stepniewska I, Kaas JH (1997) Architectonic subdivisions of the inferior pulvinar in New World and Old World monkeys. Visual Neuroscience 14:1043–1060.

Stepniewska I, Cerkevich CM, Kaas JH (2015) Cortical Connections of the Caudal Portion of Posterior Parietal Cortex in Prosimian Galagos. Cerebral Cortex 26:2753–2777.

Takakuwa N, Isa K, Onoe H, Takahashi J, Isa T (2021) Contribution of the Pulvinar and Lateral Geniculate Nucleus to the Control of Visually Guided Saccades in Blindsight Monkeys. The Journal of Neuroscience 41:1755–1768.

Togo M, Lyu D, Huang W, Pantis S, Fisher R, Matsumoto R, Buch V, Parvizi J (2025) Electrophysiological connections linking medial pulvinar, anterior nuclei of the thalamus and the hippocampus. Brain 148:4315–4324.

Ungerleider LG, Haxby JV (1994) ‘What’ and ‘where’ in the human brain. Current Opinion in Neurobiology 4:157–165.

Vann SD, Aggleton JP, Maguire EA (2009) What does the retrosplenial cortex do? Nature Reviews Neuroscience 10:792–802.

Vesia M, Crawford JD (2012) Specialization of reach function in human posterior parietal cortex. Experimental Brain Research 221:1–18.

Wang Q, Stepniewska I, Kaas JH (2023) Thalamic connections of the caudal part of the posterior parietal cortex differ from the rostral part in galagos (Otolemur garnettii). Journal of Comparative Neurology 531:1752–1771.

Warner CE, Goldshmit Y, Bourne JA (2010) Retinal afferents synapse with relay cells targeting the middle temporal area in the pulvinar and lateral geniculate nuclei. Frontiers in Neuroanatomy Volume 4 -2010.

Webster MJ, Bachevalier J, Ungerleider LG (1993) Subcortical connections of inferior temporal areas TE and TEO in macaque monkeys. Journal of Comparative Neurology 335:73–91.

Wilke M, Turchi J, Smith K, Mishkin M, Leopold DA (2010) Pulvinar Inactivation Disrupts Selection of Movement Plans. The Journal of Neuroscience 30:8650–8659.

Yeterian EH, Pandya DN (1985) Corticothalamic connections of the posterior parietal cortex in the rhesus monkey. Journal of Comparative Neurology 237:408–426.

Yeterian EH, Pandya DN (1997) Corticothalamic connections of extrastriate visual areas in rhesus monkeys. Journal of Comparative Neurology 378:562–585.

Yu H-H, Chaplin Tristan A, Davies Amanda J, Verma R, Rosa Marcello GP (2012) A Specialized Area in Limbic Cortex for Fast Analysis of Peripheral Vision. Current Biology 22:1351–1357.

